# Phosphorus availability and arbuscular mycorrhizal fungi limit soil C cycling and influence plant responses to elevated CO_2_ conditions

**DOI:** 10.1101/2021.07.16.452715

**Authors:** L. Castañeda-Gómez, J.R. Powell, E. Pendall, Y. Carrillo

## Abstract

Enhanced soil organic matter (SOM) decomposition and organic phosphorus (P) cycling may help sustain plant productivity under elevated CO_2_ (eCO_2_) and P-limiting conditions. P-acquisition by arbuscular mycorrhizal (AM) fungi and their impacts on SOM decomposition may become even more relevant in these conditions. Yet, experimental evidence of the interactive effect of AM fungi and P availability influencing altered SOM cycling under eCO_2_ is scarce and the mechanisms of this control are poorly understood. Here, we performed a pot experiment manipulating P availability, AM fungal presence and atmospheric CO_2_ levels and assessed their impacts on soil C cycling and plant growth. Plants were grown in chambers with a continuous ^13^C-input that allowed differentiation between plant- and SOM-derived fractions of respired CO_2_ (R), dissolved organic C (DOC) and microbial biomass (MBC) as relevant C pools in the soil C cycle. We hypothesised that under low P availability, increases in SOM cycling may support sustained plant growth under eCO_2_ and that AM fungi would intensify this effect. We found the impacts of CO_2_ enrichment and P availability on soil C cycling were generally independent of each other with higher root biomass and slight increases in soil C cycling under eCO_2_ occurring regardless of the P treatment. Contrary to our hypotheses, soil C cycling was enhanced with P addition suggesting that low P conditions were limiting soil C cycling. eCO_2_ conditions increased the fraction of SOM-derived DOC pointing to increased SOM decomposition with eCO_2_. Finally, AM fungi increased microbial biomass under eCO_2_ conditions and low-P without enhanced soil C cycling, probably due to competitive interactions with free-living microorganisms over nutrients. Our findings in this plant-soil system suggest that, contrary to what has been reported for N-limited systems, the impacts of eCO_2_ and P availability on soil C cycling are independent of each other.

## 1. Introduction

The effect of increased atmospheric CO_2_ concentrations on plant productivity and its potential impacts on soil carbon (C) are regulated by nutrient availability (Ainsworth and Long, 2004; Finzi et al., 2011; Reich et al., 2014; Terrer et al., 2019). Higher plant productivity and increases in soil C with elevated CO_2_ (eCO_2_) have been observed when combined with nutrient addition (De Graaff et al., 2006; Dieleman et al., 2010; Hungate et al., 2009; Maroco et al., 2002; Piñeiro et al., 2017; van Groenigen et al., 2006). In the absence of external nutrient supplies, higher soil organic matter (SOM) decomposition under eCO_2_ contributes to meet plant nutrient demands with negative impacts on soil C (Carrillo et al., 2014; Drake et al., 2011; Finzi et al., 2006; Hoosbeek et al., 2004; Phillips et al., 2012; Reich et al., 2006). However, a pre-existent nutrient limitation may constrain productivity responses to eCO_2_ (Ellsworth et al., 2017; Reed et al., 2015; Reich et al., 2006), potentially preventing any changes in soil C. The current understanding about nutrient availability regulating eCO_2_ impacts on C cycling has been obtained mainly from northern hemisphere systems, where nitrogen (N) often constrains ecosystem productivity. But the impacts of eCO_2_ on soil C may be dependent on what the most limiting nutrient is (Dijkstra et al., 2013). Phosphorus-limited tropical and subtropical ecosystems are relevant C sinks that cover a vast area (Keenan et al., 2015; Pan et al., 2011; Soepadmo, 1993) and although the role of P availability regulating the impact of eCO_2_ on soil C storage is broadly recognised (Goll et al., 2012; Reed et al., 2015; Sun et al., 2017; Vitousek et al., 2010), the mechanisms of this control are poorly understood (Ellsworth et al., 2017; Norby et al., 2015; Terrer et al., 2019).

Opposite to N, soil P cycling is less coupled to soil C dynamics because P mobilisation is not necessarily linked with SOM decomposition. P cycling can be divided into inorganic and organic sub-cycles (Liu and Chen, 2008; Mullen, 2005). In the inorganic sub-cycle, mobilisation of inorganic P occurs via biochemical mechanisms without SOM mineralisation nor CO_2_ production, thus C and P cycles are not coupled. In the organic P sub-cycle, C and P cycles are more coupled and P mineralisation occurs via biological mechanisms that lead to SOM decomposition and CO_2_ production (McGill and Cole, 1981; Sharma et al., 2013; Tate, 1984). Most P is made available via inorganic P mobilisation, particularly with P sufficiency. But in low-P systems, mineral P sources are depleted and P derived from rock weathering is minimal (Goll et al., 2012; Reed et al., 2011). Therefore, the majority of P acquisition occurs via internal recycling of organic P sources (Fox et al., 2011; Johnson et al., 2003; Reed et al., 2011) involving SOM decomposition and CO_2_ production. Most of the research on the effects of eCO_2_ on soil P availability are focused on its impacts on inorganic P cycling via biochemical P mobilisation mechanisms. For example, higher microbial P enzymatic activity is reported with eCO_2_ (Hasegawa et al., 2016; Kelley et al., 2011; Moorhead and Linkins, 1997; Souza et al., 2017) as well as higher production of organic acids and siderophores involved in P mobilisation (Fransson and Johansson, 2010; Högy et al., 2010; Tarnawski and Aragno, 2006; Watt and Evans, 1999). Moreover, increases in fine root production and changes in pH and soil moisture with eCO_2_ can indirectly enhance inorganic P cycling in soils (Dijkstra et al., 2012; Hasegawa et al., 2016; Jin et al., 2015). It is likely that organic P cycling is also affected by eCO_2_ due to the influence of eCO_2_ on ecosystem stoichiometry (Gifford et al., 2000; Loladze, 2014, 2002) but experimental evidence for this is scarce. A better understanding about how eCO_2_ and P availability influence soil C cycling is important to better predict changes in global soil C due to eCO_2_ (Finzi et al., 2011; Goll et al., 2012; Yang et al., 2016).

In low-P conditions, organic P cycling may help sustain plant productivity under eCO_2_ at the expense of higher SOM decomposition. A study by Jing et al, (2017) demonstrated that inputs of labile C led to higher SOM decomposition in low-P soils mediated by altered microbial community composition and increases in microbial biomass. Thus, as C allocation belowground increases with eCO_2_ (Canadell et al., 1995; Cotrufo and Gorissen, 1997; Pausch and Kuzyakov, 2018), the higher labile C availability, increased microbial biomass and activity and higher SOM decomposition may allow for a sustained nutrient release for plant uptake and growth in low-P soils. However, evidence from field experiments exposed to eCO_2_ suggests that under P-limiting conditions, changes in soil C cycling are minor or undetected (Castañeda-Gómez et al., 2020; Dijkstra et al., 2013; Drake et al., 2016) and that the CO_2_ fertilization effect is reduced or not observed (Conroy et al., 1992; Deng et al., 2015; Ellsworth et al., 2017; Goll et al., 2012; Peñuelas et al., 2012; Zhang et al., 2014). Thus, if P availability becomes limiting, ecosystem productivity and microbial-mediated SOM decomposition may be hampered, preventing further changes in soil C cycling. Soil microbial communities play a key role in SOM decomposition and both biological and biochemical P mineralisation and mobilisation. Moreover, the responses of SOM decomposition and C stabilisation to P availability have been found to be largely mediated by changes in saprotrophic microbial community composition and activity (Fang et al., 2019; Feng and Zhu, 2019; Liu et al., 2018; Soong et al., 2018; Yuan et al., 2021) and by symbiotic fungi such as arbuscular mycorrhizal (AM) fungi (Xu et al., 2018). Yet the role of saprotrophic and mycorrhizal fungi altering SOM decomposition under eCO_2_ conditions and low-P availability is poorly understood.

AM fungi are known to greatly contribute to P mobilisation under eCO_2_ conditions, especially in ecosystems with low soil P availability (Delucia et al., 1997; Matamala and Schlesinger, 2000; Terrer et al., 2016) where AM fungi tend to be more abundant (Treseder and Cross, 2006). AM fungi also alleviate P limitation conditions for plants (Soka and Ritchie, 2014; Willis et al., 2013) possibly via excretion of P hydrolytic enzymes (Joner et al., 2000; Sato et al., 2015) and certainly via uptake of inorganic nutrients from the soil solution (Javot et al., 2007; Smith et al., 2011). They have also been linked with increased SOM decomposition and changes in soil C cycling with climate change (Carrillo et al., 2015; Cheng et al., 2012; Talbot et al., 2008; Wei et al., 2019). Higher C transfer to AM fungi under eCO_2_ enhances AM fungi growth and activity (Mohan et al., 2014) and so their contribution to plant P acquisition in P-limited ecosystems may be enhanced with eCO_2_ (Cavagnaro et al., 2011). On the other hand, AM fungi can also suppress eCO_2_-mediated increases in plant growth due to increased P immobilisation in AM fungal biomass and increased competition over P with saprotrophic communities (Jin et al., 2015) that can simultaneously decrease AM fungi-mediated SOM decomposition (Talbot et al., 2008). If P availability is low, the role of AM fungi mobilising P and promoting plant growth may be constrained due to P-limitation (Treseder and Allen, 2002), which might further regulate impacts of eCO_2_ on ecosystem productivity in P-limited ecosystems. Due to the significant role of AM fungi on P and SOM dynamics, predictions of altered soil C stocks under eCO_2_ require an understanding how eCO_2_ can change AM fungal activity, their impact on saprotrophic communities and plant responses to eCO_2_, particularly under P limitation.

In this study, we explored the role of P availability and AM fungi mediating responses of SOM decomposition and plant productivity to eCO_2_. We did this by manipulating P availability, AM fungal presence and atmospheric CO_2_ levels in a pot experiment. An Australian native grass species (*Microlaena stipoides*) with the ability to grow in a wide range of P availability conditions and forming arbuscular mycorrhizal associations (Clark et al., 2014; Hill et al., 2010) was grown in chambers that allowed the continuous isotopic labelling of plant tissues so we could directly assess changes in SOM-derived C in different C pools: respired CO_2_ (R; the product of decomposition), dissolved organic C (DOC; considered an active pool of C) and microbial biomass C (MBC; the agent of SOM decomposition). This approach allowed for a holistic assessment of the impact of eCO_2_, P availability and AM fungal presence on soil C cycling and plant growth. Changes in saprotrophic communities were also measured (using phospholipid fatty acids) in relation to eCO_2_ conditions, P treatment and AM fungal presence. These responses, along with nutrient contents in plant tissues and soil extracts (dissolved nutrients), were collected to enhance result interpretations.

We hypothesised that for plants grown under eCO_2_ and enhanced P availability – achieved by P addition – a fertilisation effect would be observed as increased plant biomass without changes in SOM decomposition (**Figure 1A**) since P addition will allow for plant requirements to be met under eCO_2_. Contrarily, if P is low, an eCO_2_ fertilisation effect would only occur with increases in SOM decomposition, evidenced as higher SOM-derived respired CO_2_, MB and MB (**Figure 1B**). Alternatively, we hypothesised that no fertilisation effect would be observed if soil C and P cycling are decoupled in this experimental system (**Figure 1C**). The presence of AM fungi would increase SOM decomposition as a mechanism to obtain P from SOM via changes in saprotrophic communities. This effect will be observed in the different C pools, particularly with low P conditions and eCO_2_ since both low P availability and eCO_2_ promote AM fungal colonisation rates and activity.

**Figure 1.**
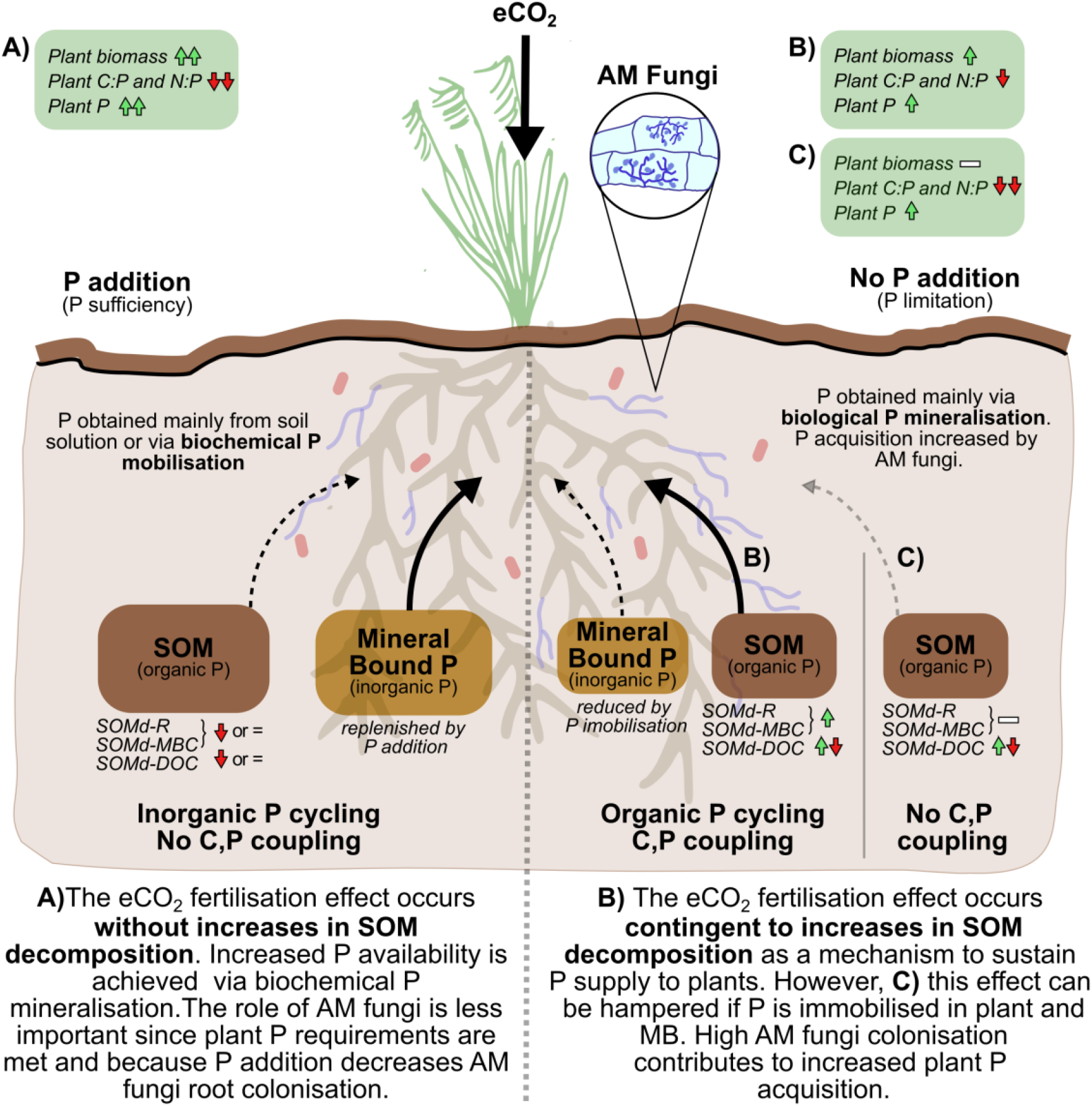
Conceptual framework and expected responses of measured variables in the experiment (green arrow: increase; red arrow: decrease; -: no effect). Increases in plant biomass under eCO_2_ conditions are dependent on nutrient availability. In low phosphorus (P) soils, **A)** P addition (left side) will support an increase in plant biomass with eCO_2_ (fertilisation effect) without major impacts in soil organic matter-derived (SOMd) C pools: R, respired CO_2_; MB, microbial biomass; DOC, dissolved organic C. Without P addition (right side), **B)** P acquisition may be more reliant on organic P cycling (coupled C and P cycles) with enhancements in plant biomass with eCO_2_ only observed if SOM decomposition and biological P mineralisation increase. Immobilisation of P by plants, AM fungi and microbial biomass becomes relevant in this scenario further increasing SOM decomposition **C)** If P availability is too low, becoming limiting, increased P immobilisation and higher P competition will decrease the impact on SOM decomposition and plant biomass.

## 2. Materials and methods

We set up a factorial experiment with three factors: **phosphorus (P) treatment**: P addition (+P)/control; **AM fungi**: Active AM inoculum (AM)/Non-mycorrhizal (Sterile AM inoculum, NM) and **CO_2_**: ambient (400 ppm - aCO_2_) and elevated (640 ppm - eCO_2_) for a total of 8 treatment combinations with 4 replicates each (+P/AM/aCO_2_; +P/AM/eCO_2_; +P/NM/aCO_2_; +P/NM/eCO_2_; control/AM/aCO_2_; control /AM/eCO_2_; control /NM/aCO_2_; control /NM/eCO_2_; n=32, **Figure SI 1A**). Plants of *Microlaena stipoides* (Labill.) R.Br., a native Australian C_3_ grass with a broad P range, were grown from seed in 125mm pots containing around 500 g of dry soil. Plants were grown in growth chambers for twelve weeks and soil water content was kept between 15 and 20 % gravimetric content by addition of MilliQ water as needed every 2-3 days.

### 2.1. Soil characteristics and treatment preparation

Soil was collected from a Cumberland plain natural woodland in Western Sydney (33°37’01” S, 150°44’26” E, 20 m.a.s.l) near the *Eucalyptus* Free-Air CO_2_ experiment (EucFACE). Soil at this site is an aeric podsol, slightly acidic (pH 5.38 ± 0.02 at 0-10 cm) (Ross et al., 2020), with a N content of 677 mg Kg^-1^, low total C (1.8%, 0-15 cm), and P content (76.28 mg Kg^-1^) (Hasegawa et al., 2016). Previous studies have demonstrated that both the vegetation (Crous et al., 2015) and soil fauna (Nielsen et al., 2015) are limited by P within this ecosystem. Soil was sieved (to 2 mm) and sterilised (gamma irradiated, 50 kGy) to remove viable AM fungal propagules. To reintroduce a homogenous microbial community to all pots, excluding AM fungi, we prepared a microbial inoculum. For this, approx. 3 Kg of freshly collected soil from a nearby grassland was mixed with water in a 1:3 proportion by volume and the suspension passed through a 20 μm mesh sieve to remove AM fungal spores (Brundett and Australian Centre for International Agricultural Research, 1995). Prior to potting, the filtrate was added to the sterile soil at a rate of 50 mL filtrate per Kg soil. The soil was incubated for a week at room temperature, mixing it daily.

The soil was then divided into four subsets based on whether they were to receive additional P and AM fungal inoculum. For the former, triple super phosphate (Richgro, super phosphate fertiliser supplement 9.1% P w/w) was added in a rate of 0.4 g/kg dry soil. The triple super phosphate powder was weighed, diluted and sprayed into the soil while low P treatments received MilliQ water only. The soil was then thoroughly mixed and left to settle for a day. Next, AM fungi treatments were created by applying AM fungal inoculum in a 1:10 AM fungi inoculum:soil with non-mycorrhizal controls produced by autoclaving the live AM inoculum (121 °C, two hours) and applying it to the soil in the same manner (**See supplementary information for AM inoculum production**). Once the four types of soils were prepared, soil was added to pots and five surface sterilised *M. stipoides* seeds (30 % H_2_O_2_ for 10 minutes followed washing) were sown per pot. Pots were randomly split and placed in aCO_2_ and eCO_2_ chambers. After two weeks of growth, pots were thinned to one plant per pot. Unplanted pots (n=16) with the different P and CO_2_ treatments (+P/NM/aCO_2_; +P/NM/eCO_2_; control/NM/aCO_2_; control/NM/eCO_2_ n=4 each), were included to estimate effects on SOM decomposition in the absence of plants and AM fungi (**Figure SI 1A**). Unplanted pots were all prepared using soil with sterile AM inoculum and were kept under the same conditions as the planted pots for the duration of the experiment.

### 2.2. Growth Chambers set up

The growth chambers (six in total, three per CO_2_ treatment) were modified using the approach of Cheng and Dijkstra (2007) to achieve a continuous ^13^C- labelling of plant tissues in both aCO_2_ and eCO_2_ treatments. The chambers were adapted to take an influx of naturally ^13^C-depleted CO_2_ (δ^13^C= −31.7 ‰ ±1.2) delivered during the photoperiod, combined with a scrubbing system made of a soda lime-filled PVC tube (72 L) that allowed continuous supply of CO_2_-free air (**Figure SI 1B**). Chambers were adjusted to a 16h/8h photoperiod with 25 °C/18 °C, 60 % relative humidity, and light intensity of 900 μmol/m^2^s^1^. The chamber atmosphere was sampled frequently to confirm depletion in ^13^C. Air samples from the chambes were extracted via a pump system into a tedlar bag (Tedlar^®^ SCV Gas Sampling Bag) and analysed for δ^13^C in a PICARRO G2201i isotopic CO_2_/CH_4_ analyser (Picarro Inc., Santa Clara, CA, USA). To avoid plant-uptake of ^13^C from outside the chambers, watering and all other manipulations were performed during the night period aided by green light.

### 2.3. Harvest and sample processing

#### Gas sampling of the plant-soil system

After twelve weeks of growth, we quantified rates of total respired CO_2_ (R) and its C isotopic composition as described by Carrillo et al., (2015, 2014). Briefly, pots were placed on an elevated platform inside a water filled tray and covered with a PVC chamber (45 cm H x 15 cm D) adapted with an air-tight rubber stopper for air sampling. Free-CO_2_ air was circulated for 2 hours using an aquarium pump connected to a CO_2_ scrubber (50 cm H x 4 cm D PVC tubing filled with soda lime). After 2 hours of scrubbing, an air sample was taken to determine baseline CO_2_ concentrations using a 7890A gas chromatograph with a G1888 network headspace sampler (Agilent Technologies, USA). Later, the CO_2_ scrubbers were removed and the pump reconnected to the PVC chamber to allow for air circulation while pots were incubated. After two to three hours of incubation, a gas sample was collected in an airtight gas collection bag (Tedlar^®^ PVDF, 1L) using an aquarium pump system and analysed for its C isotopic composition.

#### Plant and soil harvest

One day after gas sampling, pots were destructively harvested. Aboveground biomass was cut and roots separated from the soil and washed. Plant aboveground biomass, roots and a subsample of fresh soil were oven dried at 60 °C for measurements of dry plant biomass, soil gravimetric water content and total nutrients. A fresh soil subsample was stored at −20 °C for microbial community analyses and the remaining fresh soil was used for assessments of dissolved nutrients and microbial biomass.

#### Plant, soil nutrients and microbial biomass C and N

Two sub-samples of soil were weighed. Dissolved nutrients in soil were extracted from the first subsample with a 0.05M K_2_SO_4_ solution in a 4:1 solution to soil ratio, shaking at 180 rpm for an hour. Samples were filtered through a Whatman # 42 filter paper and frozen (-20°C) until analyses. The second subsample was fumigated for 5 days with chloroform and then nutrients were extracted as for unfumigated samples. The fumigated and unfumigated extracts were analysed for total dissolved organic C and N (Shimadzu^®^ TN, TOC-L, Japan) and microbial biomass C and N calculated by subtracting fumigated and unfumigated samples (Vance et al., 1987). The volume left of these K_2_SO_4_ extractions was used to obtain the isotopic composition of DOC and microbial biomass (see section below). Phosphates were extracted following the Bray 1-P method for acidic soils, in a 1:7 solution:soil ratio using a 0.03M NH_4_F solution in 0.025 M hydrochloride (HCl) adjusted pH to 2.6 ± 0.05 with HCl (Rayment et al., 2010). Samples were manually shaken for 60s and immediately poured over a Whatman # 42 filter paper. Collected extracts were analysed by colorimetry (AQ2 Discrete Analyser, SEAL Analytical, Mequon, WI, USA).

Total P and N were determined from ground oven-dried soil and root samples. Total P concentration was analysed by an X-ray fluorescence spectrometer (PANalytical, εpsilon 3. 10Kv, 0.9mA. Lelyweg, Almelo, Netherlands) while total C and N from soil, roots and soil extracts were analysed along their isotopic composition.

### 2.4. Respired, dissolved organic C (DOC) and microbial biomass isotopic composition and partitioning

Gas samples from plant-soil system incubations were analysed in a PICARRO analyser (G2201i; Picarro, Santa Clara, CA, USA. Precision values below 0.16‰) for the δ^13^C and CO_2_ concentration of the total respired CO_2_ (R). For the isotopic composition of the DOC and microbial biomass C (MBC), fumigated (*f* and unfumigated (*uf*) soil K_2_SO_4_ extracts were oven dried at 60°C. The dried extracts were scraped and weighed for analysis on a Thermo GC-C-IRMS system (Trace GC Ultra gas chromatograph, Thermo Electron Corp., Milan, Italy; coupled to a Delta V Advantage isotope ratio mass spectrometer through a GC/C-III); University of California, Davis Campus, USA). The δ^13^C of unfumigated soil extracts was used as the isotopic composition of the DOC while δ^13^C of MBC (δ^13^C_MBC_) were calculated from δ^13^C values of both fumigated (*f* and unfumigated (*uf*) extractions (**See supplementary information for calculations of the isotopic composition of the microbial biomass**). Samples of dried soil were also analysed on the Thermo GC-C-IRMS system for C, N and δ^13^C.

To calculate the fractions of SOM-derived C in the total respired CO_2_ (R), dissolved organic C (DOC) and microbial biomass C (MBC) we used isotopic partitioning as: SOM.C*_R,DOC,MBC_* = (δ^13^C*_R,DOC,MBC_* - δ^13^C_*p*_) / (δ^13^C_*control*_ - δ^13^C_*p*_). Where δ^13^C*_R,DOC,MBC_* were the δ^13^C of either the CO_2_ measured in the total respired CO_2_ (R), dissolved organic C (DOC) or microbial biomass C (MBC)) from each pot; δ^13^C_*p*_ is the isotopic ratio of the plant source averaged across P treatments per CO_2_ condition (root biomass at aCO_2_ δ^13^C= −40.01±0.08 and eCO_2_ δ^13^C= −43.20±0.04) and δ^13^C*_control_* is the average δ^13^C of the native SOM source obtained from the dried bulk soil across CO_2_ and P conditions (δ^13^C= −25.31±0.02). The fractions of plant-derived C were obtained by subtracting SOM-derived fractions from the unit. Measured total respired CO_2_ rates, DOC and MBC were partitioned into SOM-derived and plant-derived, using these fractions to obtain the mass of the SOM derived R, MBC and DOC pools. In this study, we focus only on the SOM-derived C pools and fractions.

### 2.5. Microbial community analysis: Phospholipid-derived fatty acids (PLFA) and Neutral lipid-derived fatty acids (NLFA)

Soil PLFAs were extracted to assess the overall microbial communities while the 16:1ω5c NLFA was used as an indicator of arbuscular mycorrhizal (AM) fungi presence and abundance (Olsson, 1999). Freeze-dried soil from planted pots (n=32) were extracted following the protocol by Buyer & Sasser (2012) with modifications by Castañeda-Gómez et al. (2020) (**See supplementary information for PLFA and NLFA analyses**). Functional groups were defined as shown in **Table SI 1**. Fungal to bacterial ratio (F:B) was calculated by dividing fungal PLFAs (not including AM fungi) by the sum of bacterial PLFAs. The sum of individual lipids was used as an indicator of the size of the community (μg PLFA g^-1^ dry soil).

### 2.6. Statistical analyses

The effect of CO_2_ condition, AM fungi treatment and P addition and their interactions on the response variables was analysed with a linear model fitted with the function “lm” from the stats package in R version 3.3.2 (R Core Team 2016). This approach was used instead of a mixed effects modelling approach since pots were moved among chambers with the same CO_2_ treatment so it was not possible to estimate a random effect associated with each chamber. The normality of the residuals of each model was inspected to check the appropriateness of the fit and transformations were performed when needed. Statistical significance was determined performing an ANOVA (Analyses of variance) with the “Anova” function (“car” package, Fox et al. 2017). Multiple mean comparisons were performed using the Tukey test using the “glht” function (“multcomp” package, Hothorn et al. 2017). As soil moisture can affect soil respiration measurements, we tested for correlations between soil moisture and the total respired CO_2_, SOMd- respired CO_2_ fraction and total mass of SOMd- respired CO_2_. For response variables with a significant correlation with moisture, a 3-way ANCOVA was performed with soil moisture as covariate, to account for the variability brought by the slightly different moisture contents at the time of CO_2_ sampling. The homogeneity of the regression slopes, normality of residuals and homogeneity of variances was tested when performing the ANCOVA. Significance levels were: ≤ 0.1(.), ≤ 0.05(*), ≤0.01 (**), ≤ 0.001(***).

Microbial communities were analysed as the PLFA-based total microbial biomass (sum of all individual lipids, in μg PLFA g^-1^ dry soil), as absolute microbial abundance (reflecting the size or biomass of the community, in μg PLFA g^-1^ dry soil) and as relative abundance (in percentage, reflecting microbial community composition) of the different microbial groups and as individual lipids. To visualise significant three-way interactions of experimental factors in the abundance of microbial groups from the ANOVA, the “emmip” function from the “emmeans” package (Lenth et al., 2020) was used to show the estimated marginal means from the fitted linear model.

## 3. Results

### 3.1. Influence of elevated CO_2_, P availability and AM fungi on SOM decomposition and C pools

Higher SOM-C losses under eCO_2_ were expected for control P conditions (low P) and particularly when AM fungi were inoculated **(Figure 1)**. However, we found that in general, there was not a significant interactive impact of the three experimental factors on the respired CO_2_, microbial biomass C (MBC) and dissolved organic C (DOC) as initially expected. Instead, we found that the interaction of AM and P treatments was relevant determining the fate of these soil C pools, regardless of eCO_2_. AM and P treatments significantly affected the total respired CO_2_ (**Figure 2**), which decreased with P addition but only for the NM treatment (P<0.05, Tukey’s multiple comparison). However, we did not find any significant effects of CO_2_ conditions, AM or P treatments on the SOM-derived respired CO_2_ **(Figure 2)**. On the other hand, most of the microbial biomass C (MBC) was derived from SOM (above 80%, **Table SI 2**) and while neither the total microbial biomass C nor the SOM-derived MBC (**Figure 2**) were affected by the experimental factors, the C sources used by the microbial biomass were affected by the interaction between AM and P treatment (**Table SI 2**). The fraction of MBC derived from SOM marginally increased by 6% with P addition in NM treatments (P=0.1, Tukey’s multiple comparison. **Table SI 2**). Similar to the MBC pool, most of the DOC was derived from SOM (above 80%, **Table SI 2**) and both the total DOC and SOM-derived DOC were affected by the interaction between AM and P treatments (**Figure 2**) with P addition significantly increasing the SOM-derived DOC in NM treatment (P=0.001, Tukey’s multiple comparison). Finally, although not evidenced in the total DOC or SOM-derived DOC, the fraction of DOM-derived DOC significantly increased with eCO_2_ (**Table SI 2**).

**Figure 2.**
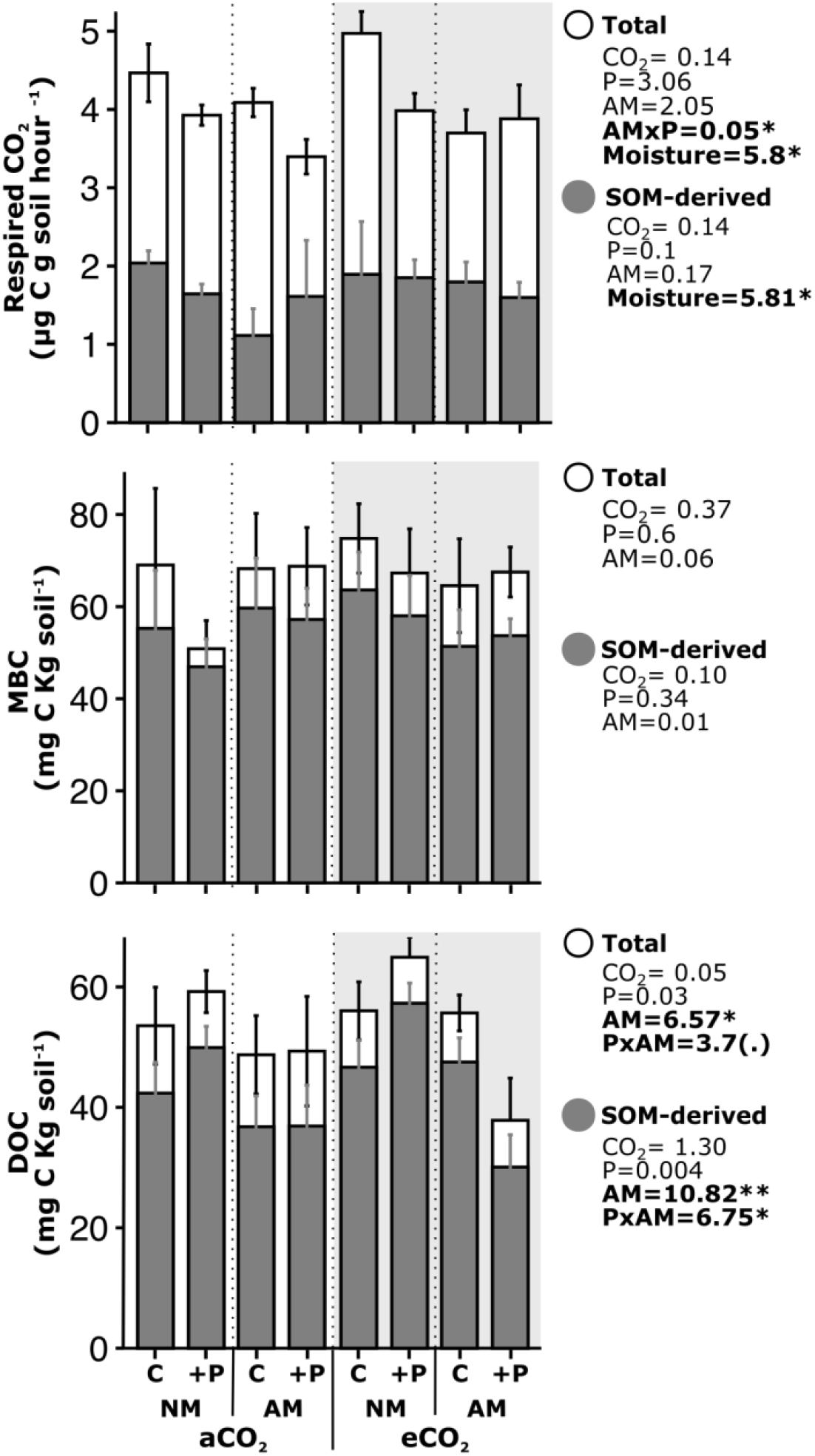
Mean values (± standard error, n=4) of the total(white) and SOM-derived (grey) respired CO_2_, microbial biomass C (MBC) and dissolved organic C (DOC) per CO_2_ condition, AM fungi and P addition treatment (C: control and +P: P addition). CO_2_ conditions are shown on light grey (elevated) and white (ambient) backgrounds. F_*(1,24)*_ values of ANCOVAs (for respired CO_2_) and ANOVAs from linear model for response variables displayed on the right. Significance levels: ≤ 0.1(.), ≤ 0.05(*), ≤0.01 (**), ≤ 0.001(***). Only significant interactions of the treatments displayed.

### 3.2. Soil nutrient responses to elevated CO_2_, P availability and AM fungi

We expected eCO_2_ to increase C and nutrient availability, P addition to increase soil P, and AM fungi to decrease soil P and N due to higher nutrient acquisition and immobilisation by the fungal biomass. These expectations were partially supported with our observations. eCO_2_ increased total soil N, but only for control P pots while dissolved N marginally decreased with eCO_2_ for AM pots (**Table 1**). Total soil P was unaffected by eCO_2_ conditions while soil dissolved P significantly decreased with eCO_2_ conditions, particularly for control P and AM pots (**Table 1**). P addition increased total soil P while AM decreased total P and dissolved N (**Table 1**). Finally, we detected lower dissolved C:N with P addition for AM pots but higher dissolved C:N for AM pots with eCO_2_ (**Figure 3**). Dissolved C:P decreased with P addition for pots under eCO_2_ conditions while it increased with eCO_2_ for AM pots (**Figure 3**).

**Table 1.**
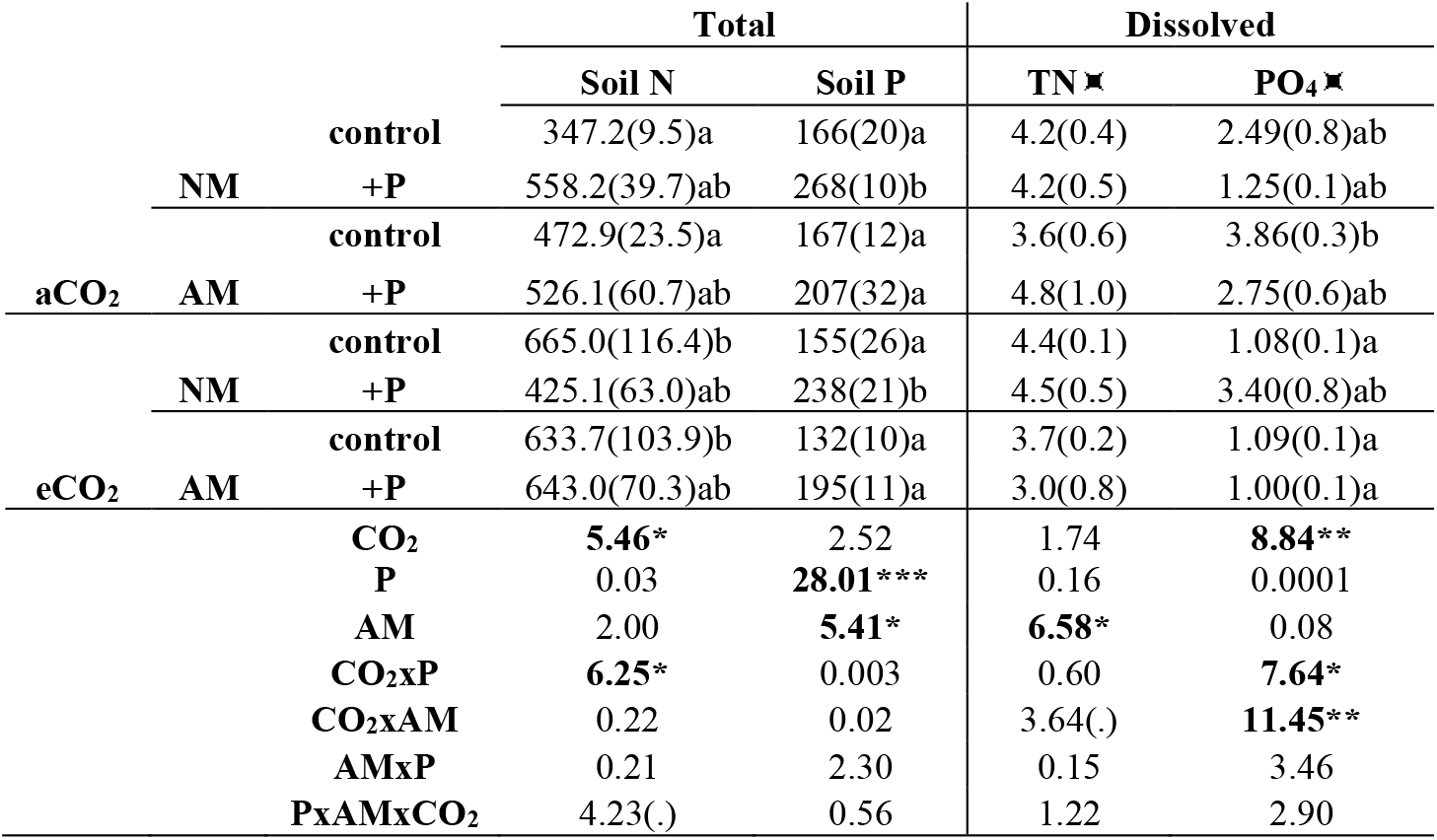
Soil nutrients (total and dissolved (ppm)) per CO_2_ condition, AM fungi and P addition treatment. Dissolved nutrients as: DOC – total dissolved organic Carbon, TN – total nitrogen and phosphates (PO_4_). Mean values (± standard error, n=4) followed by the same letter are not significantly different (p≤ 0.05, Tukey multiple comparison test), no letters indicate non-significant effect of the treatments. AM: Arbuscular mycorrhizal and NM: non-mycorrhizal. Below, ANOVA results from linear model (lm) for response variables (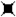 natural log transformed). F*_(1,24)_* values displayed with significance levels: ≤ 0.1(.), ≤ 0.05(*), ≤0.01 (**), ≤ 0.001(***).

**Figure 3.**
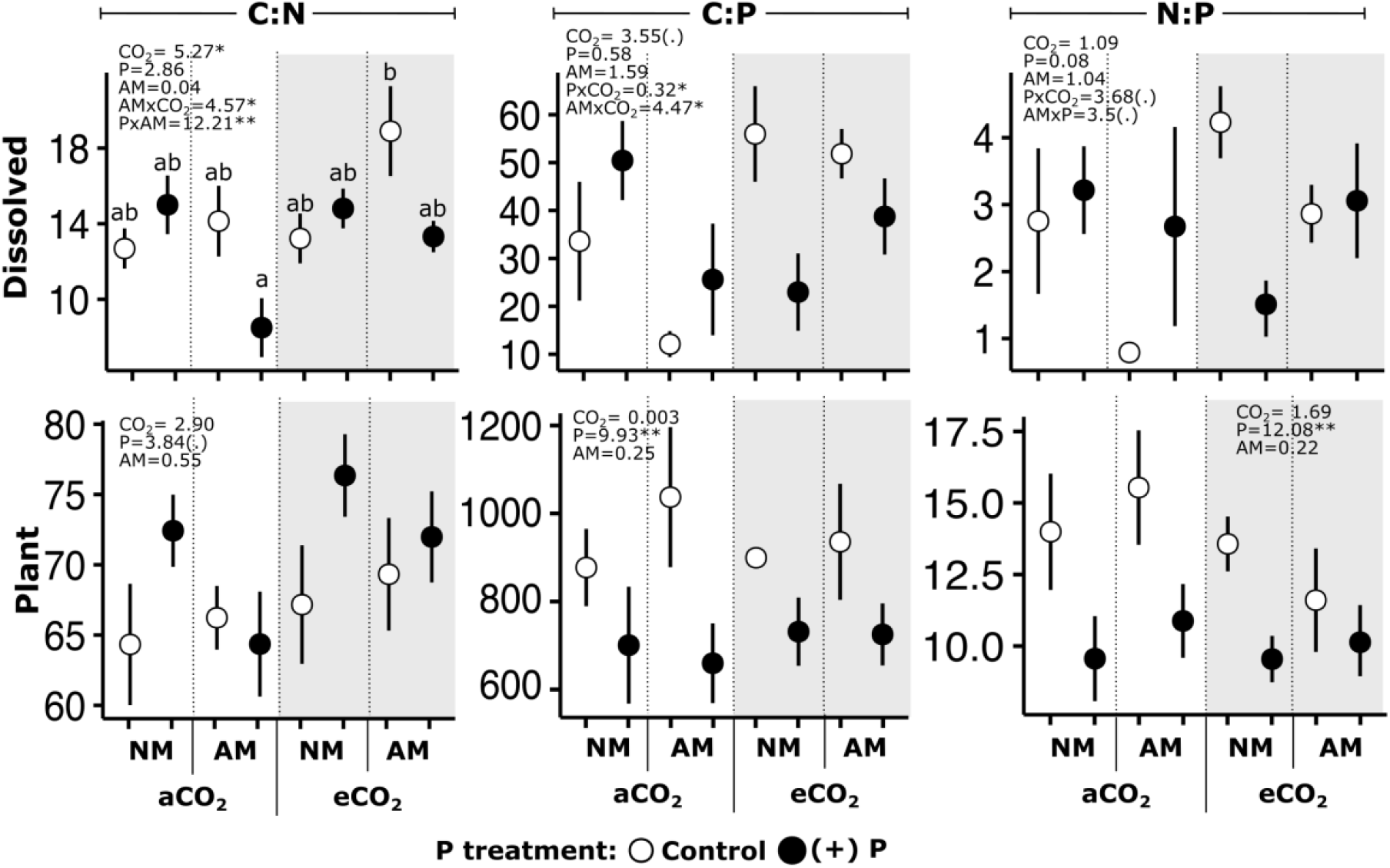
Mean values (± standard error, n=4) of mass nutrient ratios (columns) in plant and soil solution (dissolved) per treatment. eCO_2_ conditions shown as grey areas and P treatment as white(control) or black (+P) circles. NM (non-mycorrhizal). F*_(1,24)_* values of ANOVAs from linear model for response variables displayed. Significance levels: ≤ 0.1(.), ≤ 0.05(*), ≤0.01 (**), ≤ 0.001(***); only significant interactions of the treatments displayed. Treatments with different letters are significantly different (p≤ 0.05, Tukey multiple comparison test).

### 3.3. Plant responses to elevated CO_2_, P availability and AM fungi

We expected that plant biomass would increase with eCO_2_ when P availability was high, and be unresponsive to eCO_2_ when P availability was low unless AM fungi were present, supporting further plant growth by increasing SOM decomposition and P uptake (**Figure 1**). We did not observe a significant impact of eCO_2_ on aboveground plant biomass regardless of the P and AM treatments. However, eCO_2_ effects increased root biomass when AM fungi were present, regardless of the P treatment (**Table 2**). Increases in plant biomass with P addition were not observed either, but we expected to see increases in plant P concentrations in response to P addition and decreases in plant C-to-nutrient ratios with eCO_2_ as an indication of nutrient resorption and higher nutrient immobilisation, which would be particularly important in low P conditions. Supporting our predictions, we observed increases in plant P concentration as well as the plant P pool in both shoot and roots with P addition (**Table 2**). As a consequence, plant C:P and N:P significantly decreased with P addition (**Figure 3**). However, C-to-nutrient ratios in plant tissues were not strongly affected by eCO_2_ and no clear evidence for increased P immobilisation under eCO_2_ was found. On the other hand, plant N increased with AM presence but only under eCO_2_ (**Table 2**). Finally, higher plant C:N was observed with eCO_2_, whereas plant C:N increased with P addition for NM pots (**Figure 3**).

**Table 2.**
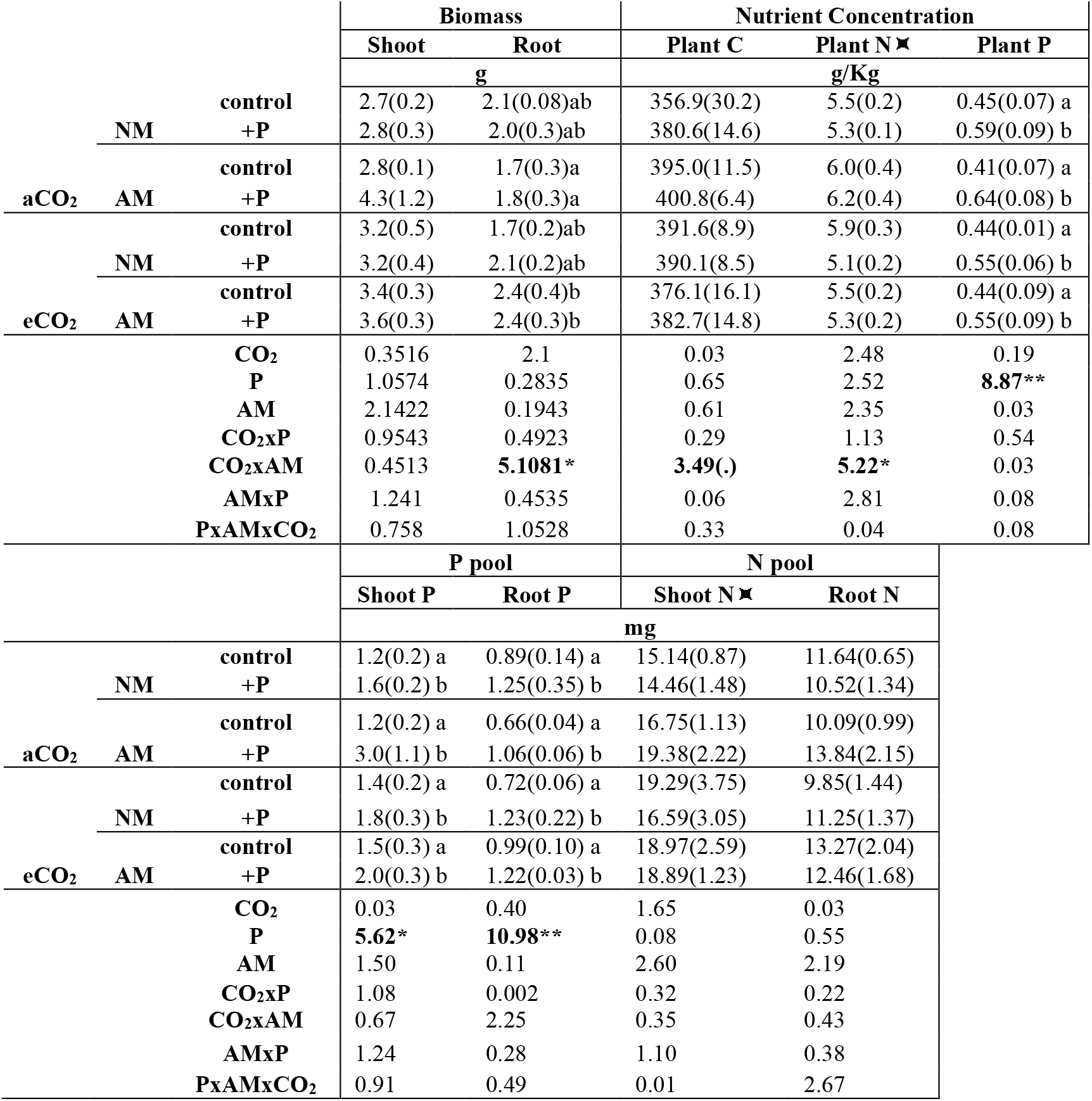
Plant biomass (shoot and roots), nutrient contents and plant P and N pools (plant P and N content per unit of shoot and root biomass) per CO_2_ condition, AM fungi and P addition treatment. NM (non-mycorrhizal). Mean values (± standard error, n=4) followed by the same letter are not significantly different (p≤ 0.05, Tukey multiple comparison test), no letters indicate non-significant effect of the treatments. Below, results of the ANOVA from linear model (lm) for response variables (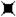 log transformed). F*_(1,24)_* values displayed with significance levels: ≤ 0.1(.), ≤ 0.05(*), ≤0.01 (**), ≤ 0.001(***).

### 3.4. Microbial communities including AM

AM fungi presence and abundance was assessed with the 16:1ω5c neutral lipid, which was high in AM-inoculated pots lacking additional P. However, P addition reduced the concentration of the AM fungal lipid in soil to a level approaching the uninoculated pots (AMxP: F*_(1,24)_*= 6.28, *P* < 0.001 **Table SI 2**). We assessed the biomass and composition of the microbial community as we expected that AM fungi would alter soil saprotrophic communities as a mechanisms to enhance SOM decomposition under low P conditions. We observed a significant increase in microbial biomass with AM fungi in low P (control pots) and eCO_2_ conditions. The increase in microbial biomass in these conditions was particularly due to the response of Gram positive bacteria and Actinobacteria (**Figure 4**). Gram negative bacteria and fungi significantly increased with eCO_2_ while protozoa increased with eCO_2_ only for NM pots **(Figure SI 2)**.

**Figure 4.**
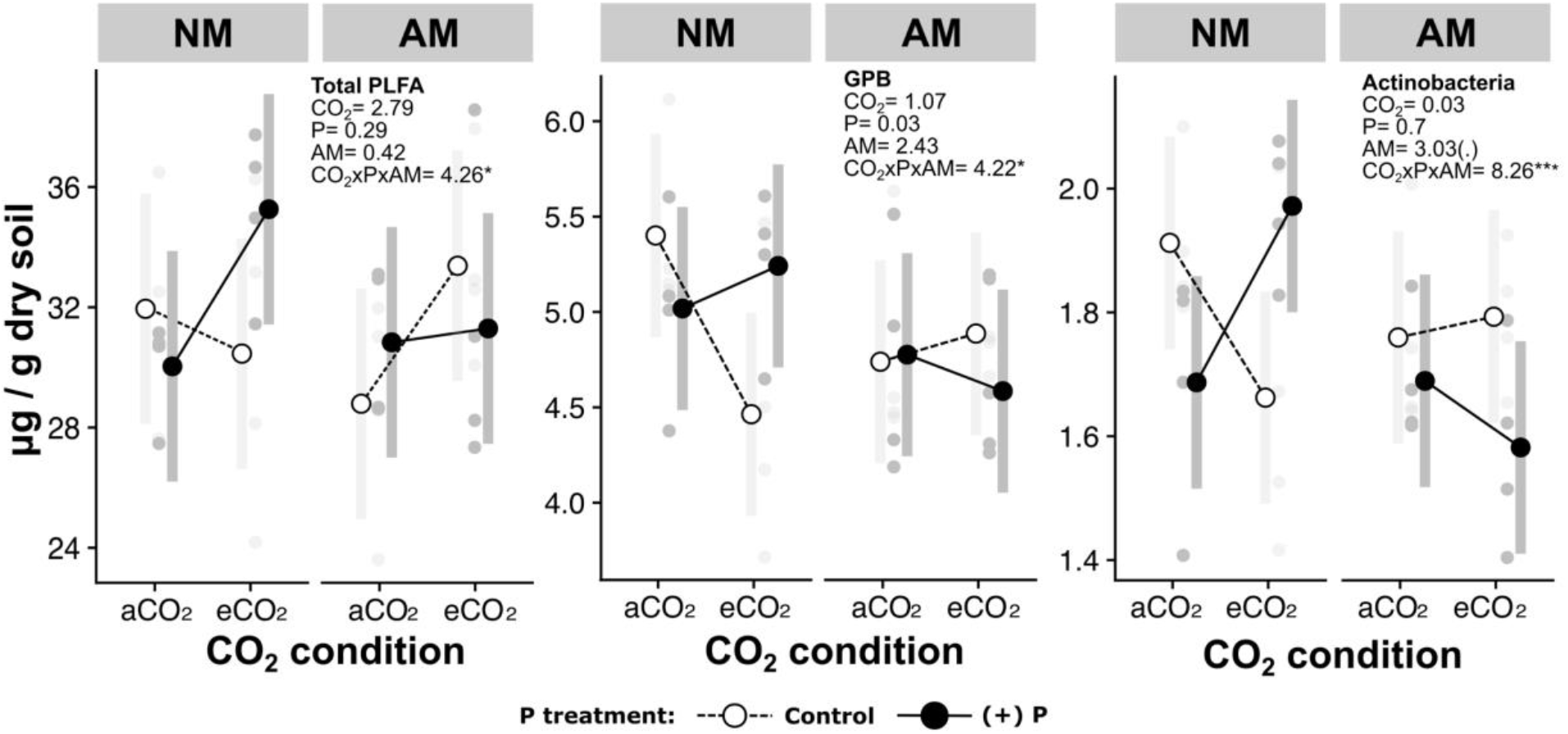
Estimated marginal means with original data (grey circles) and confidence intervals (grey bars) of the total microbial biomass, GPB (Gram positive bacteria) and actinobacteria (μg /g dry soil) from fitted lineal models per experimental treatments. P treatment as white circles and dashed lines (control) or black circles and solid lines (+P) for estimated means, and as light grey (control) and dark grey (+P) for original data. NM: non-mycorrhizal. ANOVA results displayed with significance levels: ≤ 0.1(.), ≤ 0.05(*), ≤0.01 (**), ≤ 0.001(***); only significant interactions of the treatments displayed. Grey circles are the original data input for the models.

## 4. Discussion

Our experiment investigated whether, under low P conditions, increased SOM decomposition and organic P cycling would allow for a sustained plant fertilisation effect with eCO_2_. We expected to observe increases in plant biomass under eCO_2_ conditions and low P availability paired with higher SOM derived C pools (SOM derived respired CO_2_, MBC, DOC), with further enhancements of SOM decomposition when AM fungi were inoculated due to the role of this symbiotic fungi on P acquisition. We did not find strong evidence suggesting that increases in soil C cycling with low P availability supported plant growth under eCO_2_. Instead, we found that the impacts of CO_2_ enrichment and P availability on plant growth and soil C cycling were generally independent of each other with increases in root biomass and soil C cycling under eCO_2_ occurring regardless of the P treatment. Contrary to our hypotheses (**Figure 1**), soil C cycling was enhanced with P addition, suggesting that low P conditions were limiting soil C cycling. On the other hand, root biomass increased with eCO_2_ conditions and AM presence, while microbial biomass increased with eCO_2_ in AM-inoculated pots and control P conditions mainly due to the response of Gram positive bacteria and Actinobacteria. Taken together, our findings in this plant-soil system demonstrate that C and P biogeochemical cycles may not become coupled to sustain an eCO_2_ fertilisation effect but instead, that low soil P will limit C cycling responses to eCO_2_ with AM potentially outcompeting saprotrophic communities for nutrients and hampering SOM decomposition.

The CO_2_ fertilization effect is modulated by the interaction of mycorrhizal associations and nutrient availability (Terrer et al., 2019, 2016; Treseder, 2004). In this study, although the increases in root biomass under eCO_2_ conditions occurred when AM fungi were present, this effect occurred regardless of P availability. The CO_2_ fertilization effect is a product of higher plant photosynthetic activity leading to excess C in plant tissues leading to increased growth. In this study, we observed a marginally higher plant C:N ratio with eCO_2_ (**Figure 3**) but no significant increases in plant biomass. The observed additional assimilated C was potentially allocated belowground, driving the detected increases in root biomass and AM fungi growth and activity (Cheng and Johnson, 1998; Mohan et al., 2014; Treseder, 2004). The presence of AM fungi has been previously reported to increase root biomass with eCO_2_ (Baslam et al., 2012; Dong et al., 2018; Zhu et al., 2016) and root biomass responses to eCO_2_ may not be necessarily affected by low P conditions (Jiang et al., 2020). It is hypothesised that the positive feedback of eCO_2_ and AM fungi on plant growth is mediated by the nutrient uptake role of these symbiotic fungi (Alberton et al., 2005). While P availability has been found to drive the CO_2_ fertilization effect on ectomycorrhizal plants, the response of arbuscular mycorrhizal plants to eCO_2_ seems to be more dependent on soil N availability (Terrer et al., 2019). Although we did not find that P availability mediated increases in root biomass with eCO_2_ and AM fungal presence, we observed that AM fungi marginally increased plant N contents in the aboveground biomass (**Table 2**), as well as decreased dissolved N and P in soil with eCO_2_ (**Table 1**) and plant C:N with P addition (**Figure 3**), all of this indicating that AM fungi might have promoted N uptake when P was added and facilitated the observed increases in root biomass.

We hypothesised that higher SOM decomposition would occur with low P availability as organic P cycling and biological P mobilisation mechanisms are crucial when mineral P sources are depleted (Reed et al., 2011). Contrary to our expectations, we found that P addition enhanced soil C cycling. Increases in SOM decomposition with P addition in P-limited systems have been previously reported (Cleveland et al., 2006) and can be explained by the higher affinity of P to mineral surfaces and the consequent release of labile C that promotes microbial activity (Mori et al., 2018). We expected that AM fungi presence would further enhance SOM decomposition in low P conditions (**Figure 1**). Labile C funnelled via AM fungi to the saprotrophic community may enhance soil microbial community activity and promote SOM decomposition (Frey, 2019). Although we observed higher microbial biomass under eCO_2_ and AM presence in control P conditions, we suggest that the lack of significant increases in SOM decomposition in these conditions may be due to higher competition between AM fungi and saprotrophic communities that might have limited enhanced soil C cycling (Zhou et al., 2019). The changes in plant C:N, soil dissolved C:N, total soil C:N with AM fungi explained above further support the idea that AM fungi might outcompete saprotrophic microbes in N and P uptake. Finally, we detected a decrease in SOM decomposition with P addition and AM-inoculated pots (**Figure 2**), but caution must be taken when interpreting this result since P addition also significantly reduced AM fungi presence and thus, it cannot be claimed that the reduction in SOM decomposition with P addition in AM pots is due to AM fungi presence.

We expected that SOM decomposition and, thus, organic P cycling would increase with eCO_2_ particularly under low P conditions as a mechanism to sustain plant nutrient demands (**Figure 1**). Our results show evidence for increased C cycling with CO_2_ enrichment, particularly for the fraction of SOM-derived DOC, but this effect was not dependent on P availability. Increases in DOC with eCO_2_ have been previously attributed to increases in C allocation belowground and higher rhizodeposition (Drake et al., 2011; Freeman et al., 2004; Lukac et al., 2003; Phillips et al., 2011) but it can also be due to increased SOM decomposition (Hagedorn et al., 2008, 2002). Our isotopic analyses allowed us to detect that increases in DOC under eCO_2_ in this system were due to enhanced SOM decomposition given the observed increase in SOM-derived DOC fraction (**Table SI 2**). The DOC made available via SOM decomposition can be either incorporated in the microbial biomass or lost via leaching. We did not observed changes in the SOM-derived MBC with eCO_2_ and thus the extra DOC was likely not incorporated in the microbial biomass but rather lost via leaching (Kindler et al., 2011). On the other hand, only marginally higher soil respiration with eCO_2_ was found in the present study which can be due to the low diffusivity of CO_2_ from soils, particularly with high soil moisture (Davidson et al., 2000; Hashimoto and Komatsu, 2006; Maier and Schack-Kirchner, 2014). In our experiment, volumetric soil water content was on average 23% for planted pots, which is slightly above the expected (12-21%) soil water holding capacity for a soil with sandy to loamy texture as the one used for this experiment (Datta et al., 2017; Easton and Bock, 2016).

We observed higher P concentrations in plant tissues but this was not accompanied by increases in biomass under any CO_2_ condition. Higher P concentration in plant tissues without increases in biomass can occur due to a P luxury consumption of plants, where higher P immobilisation is not necessarily related to increases in growth (Brar and Tolleson, 1975). Lack of responses to P addition in biomass (Nie et al., 2009) and abundance (Robinson et al., 1993) for *Microlaena stipoides* have been observed before and attributed to a nutrient accumulation strategy of this grass species, the time and frequency of fertilizer additions and the low response that Australian native grasses typically have to fertilizer application. In this study, only one addition of P as triple superphosphate was done at the start of this experiment. Australian native grasses are less responsive to fertilizer addition during establishment (Nie et al., 2009), which explains the lack of biomass effects of *M. stipoides* in this experiment. Also, triple superphosphate is highly soluble in water and becomes rapidly available for plant uptake (Mullins et al., 1995) by the first week of application (Ghosal and Chakraborty, 2012). Hence, increases in P concentrations in plant tissues occurred at the beginning of the experiment when AM fungi had not fully colonised the roots and so, their role in soil C transformations and plant P uptake were less relevant.

Current understanding of the impacts of eCO_2_ on plant productivity with low nutrient conditions have focused on N-limited ecosystems. Higher SOM decomposition under eCO_2_ when N availability is low occurs as a mechanism to sustain nutrient supply and plant growth, with mycorrhizal fungi aiding to deliver the mined nutrients to the plants. For P-limited ecosystems however, low P availability generally constrains ecosystem responses to CO_2_ enrichment (Ellsworth et al., 2017; Jiang et al., 2020; Reed et al., 2015; Reich et al., 2006) and the role of AM fungi mediating plant and soil C responses to eCO_2_ with low P availability is not fully clear. Thanks to the detailed measurement of soil C cycling as changes in total C pools, SOM-derived fractions and total mass of SOM-derived C pools, we show that the impacts of CO_2_ enrichment and P availability on plant growth and soil C cycling were independent of each other and are not likely to become coupled with eCO_2_ conditions. We also demonstrate that although AM fungi may contribute to increases in microbial biomass with eCO_2_ and low-P conditions, this effect may not translate into enhanced SOM decomposition due to increased nutrient competition that may limit saprotrophic communities. Our findings highlight that ecosystem responses to eCO_2_ with P limitation are different from those reported for N-limited systems and therefore, inferences of the behaviour of P-limited ecosystems based on current knowledge about N-limited ecosystems are not ideal. Moreover, our results also contribute to the current gap in knowledge regarding the impacts of soil C cycling with low P availability exposed to eCO_2_ conditions.

## 5. Acknowledgements

We thank Gavin McKenzie and Goran Lopaticki for their help with the chamber set up, maintenance and troubleshooting for the duration of this experiment. Thanks to Pushpinder Matta and Christopher Mitchell for their help with nutrient analyses and to Johanna Pihlblad and Johanna Wong- Bajracharya for their assistance during the harvest of this experiment and sample processing.

