## Supplementary Information for "Phosphorus availability and arbuscular mycorrhizal fungi limit soil C cycling and influence plant responses to elevated CO_2_ conditions"

### SUPPLEMENTARY METHODS

#### 1. AM inoculum production

To produce the AM inoculum, maize (Sweet corn, Zea mays L., D.T. Brown seeds) and leek (Leek Musselburgh, Allium ampeloprasum L., D.T. Brown seeds,) plants were grown in a 1:1 v/v mixture of sand:clay in association with *Glomus* sp (strain NBR 4.1, Auscott Narrabri, Isolator: McGee, P.A.). After 3 months of growth, mycorrhizal root length colonisation (Manoharachary and Kunwar 2002) was checked by microscopy, using the ink-vinegar staining protocol (Vierheilig and Piché 1998). After confirming successful root colonisation, aboveground plant biomass was removed and root systems extracted, chopped (to 1-2cm), homogenised with the original sand:clay potting mixture and applied to the soil. This inoculum mixture was low in organic matter content, thus we expected that the contribution of saprotrophic microbes in the AM inoculum was low relative to that in the added microbial filtrate.

#### 2. Calculations of the isotopic composition of the microbial biomass

The $\delta^{13}C$ values of both fumigated (*f*) and unfumigated (*uf*) dried extractions was used to calculate the isotopic composition of the microbial biomass as: $\delta^{13}C_{MB}= ( \delta^{13}C_f * C_f - \delta^{13}C_{uf} * C_{uf}) / (C_f – C_{uf})$. Where $\delta^{13}C_f$ and $\delta^{13}C_{uf}$ are the C isotopic ratios (‰) in fumigated and unfumigated samples, respectively; and $C_f$ and $C_{uf}$ the C content of the extracts (ppm).

#### 3. PLFA and NLFA analyses

Two g of freeze dried soil were extracted with 4ml of a methanol-chloroform-phosphate buffer. Lipids were separated by solid phase extraction (SPE) and both neutral (chloroform elution) and phospholipids (methanol elution) were collected, evaporated and mild alkaline methanolysis used to produce fatty acid methyl esters (FAMEs). Samples were re-suspended with a mixture of hexane and the 19:0 FAME as an internal standard (Methylnonadecanoate

≥98% (GC) Sigma-Aldrich, 20ppm) and analyses with gas chromatography (Agilent 7890A GC, Agilent Technologies, Wilmington, DE, USA). FAME profiles were identified using the MIDI PLFAD1 calibration mix and the software SHERLOCK version 6.2 (MIDI, Inc., DE, USA).

### SUPPLEMENTARY FIGURES

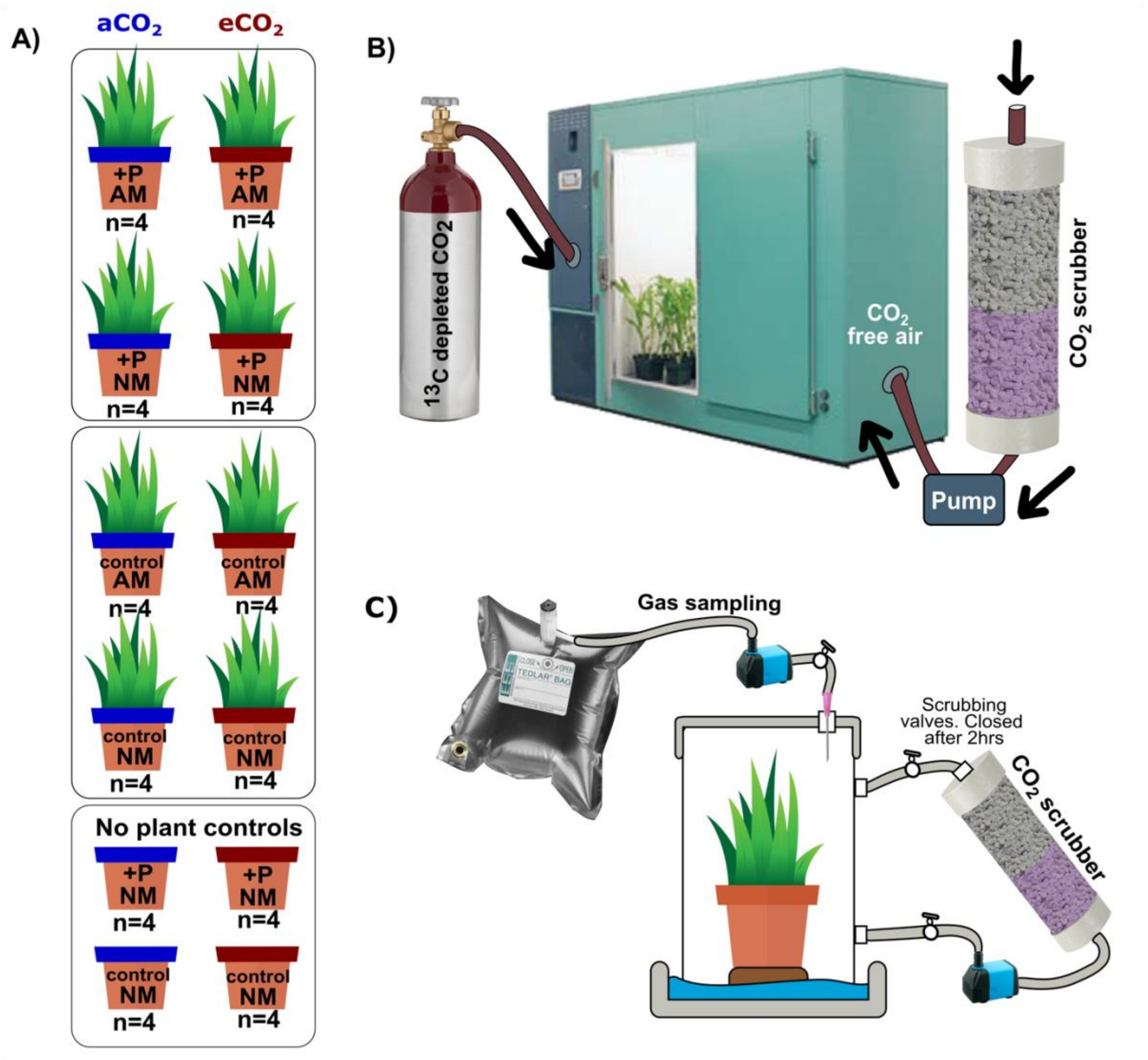


**Figure SI 1** Experimental set up including **A)** factorial design, **B)** chamber set up and **C)** gas sampling set up.

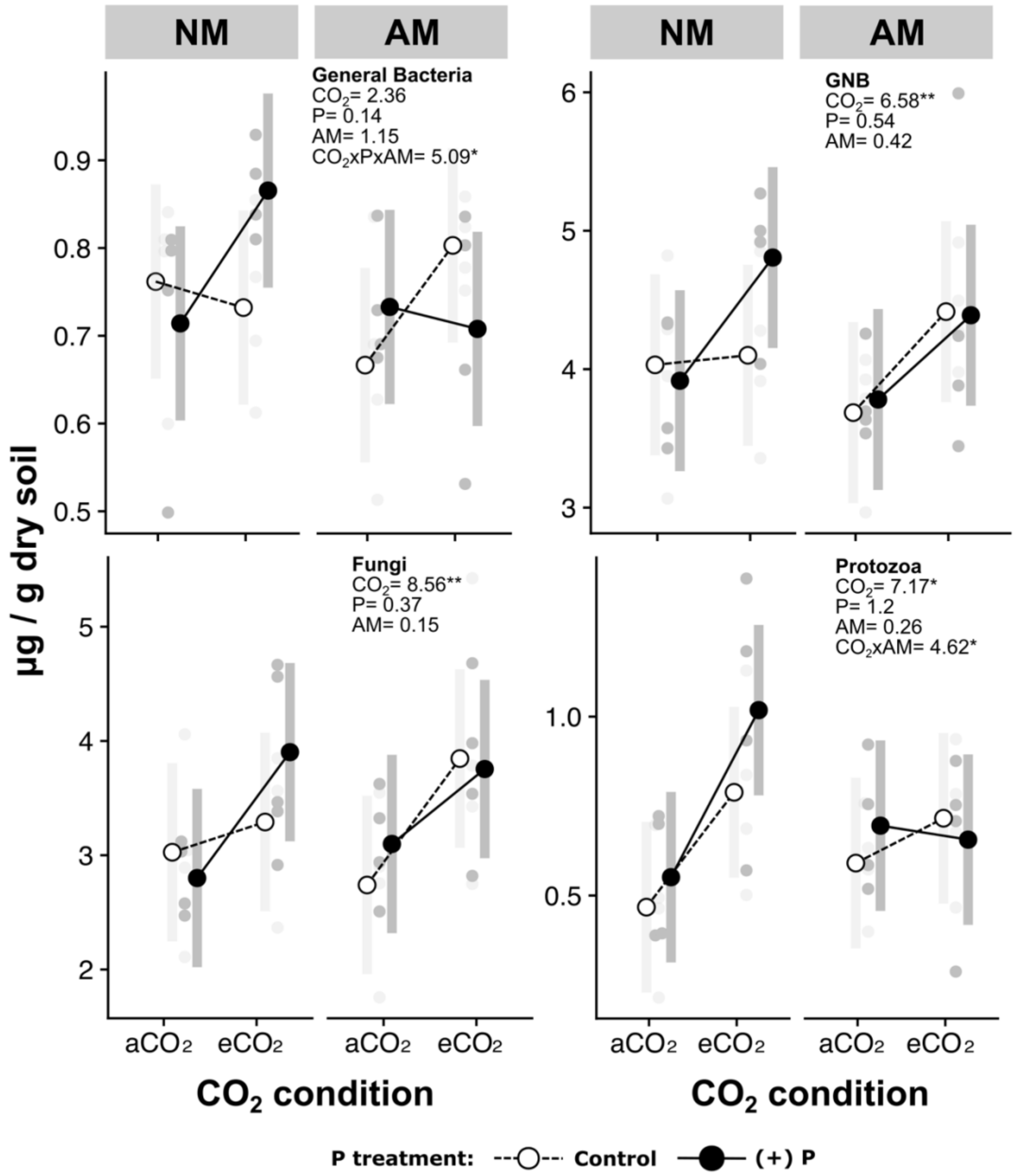


**Figure SI 2** Estimated marginal means with confidence intervals (grey bars) of the absolute abundance of the microbial groups (µg /g dry soil) from fitted lineal models per experimental treatments. P treatment as white circles and dashed lines (control) or black circles and solid lines (+P) for estimated means, and as light grey (control) and dark grey (+P) for original data. ANOVA results displayed with significance levels: ≤ 0.1(.), ≤ 0.05(*), ≤0.01 (**), ≤ 0.001(***); only significant interactions of the treatments displayed. Grey circles are the original data input for the models.

### SUPPLEMENTARY TABLES

**Table SI 1** PLFA group classification by different lipids with references using the specific lipids classified as the mentioned group.

| Group | Lipids | References |
|---|---|---|
| General bacteria | **14:0**, **15:0**, 17:0 | (Zak et al. 2000; Carrillo et al. 2011; Willers et al. 2015; Canarini et al. 2016) |
| Gram positive bacteria | **15:0iso**,**15:0anteiso**,**16:0iso**, **17:0iso**, **17:0anteiso** | (Zak et al. 2000; Bell et al. 2009; Brockett et al. 2012; Nie et al. 2013; Willers et al. 2015; Canarini et al. 2016) |
| Gram negative bacteria | **14:0 2OH**, **16:1 w7c**, 17:1 iso ω9c, **18:1ω7c**, 18:1ω5c | |
| Actinomycetes | 16:0 10-methyl, 17:1 ω 7c 10-methyl, 17:1ω7c 10-methyl, 18:0 10-methyl | (Bell et al. 2009; Carrillo et al. 2011; Brockett et al. 2012; Willers et al. 2015; Canarini et al. 2016) |
| Fungi (saprotrophic and Ectomycorrhizal fungi) | **18:2 ω6c, 18:1 ω9c** | (Frostegård and Bååth 1996; Zelles 1999; Arao 1999; Zak et al. 2000; Bell et al. 2009; Ruess and Chamberlain 2010; Brockett et al. 2012; Nie et al. 2013; Willers et al. 2015) |
| Protozoa | 20:4 ω6c, 20:5 ω3c, 20:3 ω6c | (Cavigelli et al. 1995; Canarini et al. 2016) |

Lipids in **bold** were used to calculate the fungal to bacterial ratio.

**Table SI 2** Mean values (± standard error, n=4) of the soil moisture, arbuscular mycorrhizal neutral lipid (16:1ω5c, μg NLFA g dry soil $^{-1}$) and SOM-derived fraction of respired $CO_2$, Microbial biomass C (MBC) and dissolved organic C (DOC) per $CO_2$ condition, AM fungi and P addition treatment. Below, results of the ANOVA from linear model (lm) for response variables. $F_{(1,24)}$ values displayed with significance levels: ≤ 0.1(.), ≤ 0.05(*), ≤0.01 (**), ≤ 0.001(***).

| | | | **Moisture (%)** | **NLFA 16:1ω5c** | **Respired $CO_2$** | **MBC** | **DOC** |
|---|---|---|---|---|---|---|---|
| | | **control** | 23.41(1.10) | 0.07(0.00) | 0.46(0.02) | 0.81(0.03) | 0.79(0.00) |
| | **NM** | **P** | 26.31(1.27) | 0.08(0.01) | 0.42(0.03) | 0.92(0.04) | 0.84(0.01) |
| | | **control** | 24.46(1.75) | 1.21(0.47) | 0.35(0.02) | 0.88(0.02) | 0.75(0.01) |
| **a$CO_2$** | **AM** | **P** | 21.46(1.74) | 0.06(0.03) | 0.29(0.02) | 0.84(0.05) | 0.75(0.00) |
| | | **control** | 24.79(1.66) | 0.07(0.01) | 0.57(0.10) | 0.84(0.03) | 0.83(0.01) |
| | **NM** | **P** | 24.31(1.57) | 0.06(0.01) | 0.46(0.04) | 0.86(0.02) | 0.88(0.01) |
| | | **control** | 23.72(1.14) | 1.84(0.97) | 0.51(0.10) | 0.79(0.02) | 0.85(0.03) |
| **e$CO_2$** | **AM** | **P** | 22.36(1.00) | 0.28(0.10) | 0.42(0.05) | 0.80(0.01) | 0.80(0.01) |
| | | **$CO_2$** | 0.01 | 0.61 | **7.20*** | 2.70 | 33.48*** |
| | | **P** | 0.23 | **6.33*** | 3.28(.) | 1.33 | 1.38 |
| | | **AM** | 2.84 | **8.29**** | **4.40*** | 2.50 | 25.50*** |
| | | **$CO_2$xP** | 0.19 | 0.16 | 0.34 | 0.64 | 1.53 |
| | | **$CO_2$xAM** | 0.04 | 0.64 | 0.56 | 0.95 | 2.83 |
| | | **AMxP** | 2.79 | **6.28*** | 0.02 | 4.15 (.) | 16.50*** |
| | | **PxAMx$CO_2$** | 1.54 | 0.14 | 0.09 | 2.72 | 1.60 |

†SOM-derived respired $CO_2$ fraction was not significantly correlated with soil moisture at any treatment level, so an ANOVA was performed instead of an ANCOVA

**Table SI 3** PerMANOVA results for the impact of experimental treatments on soil microbial community. Data as relative or absolute abundance and within each, as lipids belonging to groups of interests or as the individual lipids. $F_{(Df, Dfres)}$, $R^2$ and $P$ values (≤ 0.1(.), ≤ 0.05(*), ≤0.01 (**), ≤ 0.001(***)). Relative abundance as percentage and absolute abundance as μgr per gr of dry soil$^{-1}$.

| | | $CO_2$ | | | AM | | | P | | | $CO_2$ x AM x P | | |
|---|---|---|---|---|---|---|---|---|---|---|---|---|---|
| | | $F_{(1,24)}$ | $R^2$ | $P$ | $F_{(2,24)}$ | $R^2$ | $P$ | $F_{(1,24)}$ | $R^2$ | $P$ | $F_{(1,31)}$ | $R^2$ | $P$ |
| **Absolute abundance** | **Individual lipids** | 2.704 | 0.084 | **0.041*** | 0.919 | 0.028 | 0.405 | 0.509 | 0.016 | 0.743 | 2.102 | 0.065 | **0.07(.)** |
| | **Groups** | 4.715 | 0.136 | **0.019*** | 1.332 | 0.038 | 0.243 | 0.378 | 0.010 | 0.719 | 2.500 | 0.073 | **0.098(.)** |
| **Relative abundance** | **Individual lipids** | 4.803 | 0.147 | **0.004**** | 0.929 | 0.028 | 0.448 | 0.673 | 0.021 | 0.638 | 0.354 | 0.011 | 0.908 |
| | **Groups** | 9.516 | 0.265 | **0.002**** | 0.846 | 0.024 | 0.417 | 0.695 | 0.019 | 0.512 | 0.099 | 0.003 | 0.942 |
